# Genomic evidence for widespread reciprocal recognition and killing among *Pseudomonas syringae* strains

**DOI:** 10.1101/2024.06.09.598117

**Authors:** C Fautt, K.L. Hockett, S. Delattre, E. Couradeau

**Affiliations:** Department of Ecosystem Science and Management, Pennsylvania State University, University Park, Pennsylvania, United States; Department of Plant Pathology and Environmental Microbiology, Pennsylvania State University, University Park, Pennsylvania, United States; Intercollege Graduate Degree Program in Ecology, Pennsylvania State University, University Park, Pennsylvania, United States; Institute for Computational and Data Sciences, Pennsylvania State University, University Park, Pennsylvania, United States

## Abstract

Community assembly dynamics are in part driven by competition between community members. Diverse bacteria antagonize competitors through the production of toxic compounds, such as bacteriophage-derived tailocins. These toxins are highly specific in their targeting, which is determined by interactions between the tailocin’s tail fiber and competitors’ lipopolysaccharide O-antigen moieties. Tailocins play a pivotal role in mediating microbial interactions among the economically significant plant pathogens within the *Pseudomonas syringae* species complex, with the potential to alter community structure and disease progression in host plants. Previous work looking at 45 *P. syringae* strains has demonstrated that at least two phylogenetic clades of tail fibers are encoded in the conserved tailocin region across the species complex, which roughly corresponds to two clusters of targeting activity. To better understand the full diversity of tail fibers associated with tailocins in the species complex, we screened 2,161 publicly available genomes for their tailocin tail fiber content, predicted protein structures that represent the diversity of fibers, and investigated forces possibly driving the distribution of fibers throughout the species complex. Here we present evidence that while the two previously described tail fiber clades are indeed widespread among virulent *P. syringae* strains, their distribution is largely uncorrelated with phylogeny. Instead, we found that the presence of one tail fiber or the other is strongly correlated with the allelic diversity of another gene, associated with lipopolysaccharide O-antigen structure, dTDP-4-dehydrorhamnose reductase. Our findings suggest the presence of two reciprocally targeting groups of strains distributed throughout the *P. syringae* species complex that transcend phylogenetic relationships.

## Introduction

It has been estimated that the global surface area of photosynthetic leaves is approximately 10^9^ km^2^, which harbors up to 10^26^ bacteria ^1^. Of these 10^26^ bacteria, a fraction will be foliar pathogens. In a variety of crops, it has been reported that potential foliar pathogens commonly make up anywhere from 5-60% of the epiphytic community, with the pathogenic *Pseudomonas syringae* being one of the most common inhabitants ^2,3^. For potential pathogens, the successful infection of a plant depends not only on the ability of the pathogen to evade plant defenses and manipulate plant metabolism ^4^ but also to proliferate on the leaf surface, thereby increasing the chances of infiltrating the leaf apoplast ^2^. Along with adaptation to frequent fluctuations in temperature, humidity, and exposure to ultra-violet (UV) radiation, the outcome of microbe-microbe interactions on the leaf surface has been consistently shown to impact pathogenic potential, with some commensal epiphytes being able to reduce pathogenic populations and suppress disease^5–7^. Competition between pathogens and other foliar epiphytes likely plays a large role in the distribution and dispersal of pathogenic populations, with implications for the prevalence of disease, as well as the rate of gene flow, adaptation, and the evolution of novel pathogenic strains ^8,9^. Therefore, gaining a better understanding of the mechanisms plant pathogens use to compete for limited resources in the phyllosphere would not only enhance our understanding of disease ecology but also have practical implications for the management of phytopathogens.

Phytopathogenic bacteria in the *Pseudomonas syringae* species complex (PSCC) are able to kill competitors with incredible selectivity through the production of bacteriophage derived toxins called tailocins ^10,11^. Many aspects of tailed phage (*Caudoviricetes*) and tailocin targeting are similar, including a highly specific and narrow killing spectra determined by tail fiber-cell surface receptor interactions. Specifically, in one of the most well studied phages, T4, targeting and eventual infection of cells occurs through a multi-step process initiated by the binding of six long tail fibers to specific moieties in cell surface lipopolysaccharide (LPS) molecules of Gram-negative bacteria ^12^. Binding of long tail fibers is reversible, allowing the phage to ‘walk’ along the cell surface until a sufficient number of fibers are bound at the same time to trigger a conformational change in the phage baseplate that lowers the phage closer to the cell membrane ^13^. At this point, short tail fibers bind irreversibly to lipid A-inner core region of LPS, and further conformational changes in the baseplate trigger the piercing of the cell membrane ^12^. Thus, while phage tail fibers have no bactericidal activity themselves, they are essential for triggering a lethal chain reaction and ultimately determine a highly specific and narrow host range ^14,15^. Like phage, tailocins carried by PSSC are also equipped with tail fibers that are necessary for target cell recognition, and which initiate a chain reaction that leads to cell death via the forming of a pore in the cell’s membranes through which cell contents pour out ^16^ and protons flow in, disrupting the proton-gradient^17^. In contrast to bacteriophage, tailocins in PSSC carry a single set of tail fibers^11^, although tailocins in other Pseudomonads have been observed to carry multiple tail fiber genes that add functional diversity to this single set^18^.

Given the importance of tail fibers to the activity of tailocins, it is likely that they are under high selective pressure ^19^. This is true for bacteriophage, in which tail fibers show an increased rate of evolution compared to the rest of the phage genome ^19^ and there is frequent recombination of binding domains between phage families ^20^. It is likely that PSSC tailocins also experience heightened selection leading to diversification, as is evidenced by the observed recombination in the tail fiber region correlating with distinct killing spectra ^21^. It is widely believed that PSSC tailocin tail fibers bind to LPS ^22–24^, and a recent genome-wide association study expanded on this by showing that L- and D-rhamnose biosynthesis genes are highly correlated with the sensitivity of a strain to different tail fibers, suggesting that dTDP-4-dehydrorhamnose reductase (*rfbD)* is a particularly good genomic indicator of tailocin sensitivity ^25^. If rhamnose content in a cell’s LPS determines its sensitivity to tailocins, it is reasonable to hypothesize that it also imposes limitations on the tail fibers it can itself carry such that it does not self-kill. Accounting for the full genetic and structural diversity of tail fibers associated with PSSC tailocins, and investigating compatibility between LPS structure and tailocin fiber types would not only provide insights into the ecological significance of tailocins to the species complex but could also improve our ability to predict competitive outcomes between PSSC strains and suggest a robust framework for creating synthetic tail fibers capable of selectively targeting PSSC pathogens of interest ^26^.

Here, we report the results of a genetic and structural survey of tailocin-associated tail fibers found within PSSC. From a screen of 2,161 PSSC genomes, we found evidence for the circulation of three distinct recombinant tail fiber ‘types’ within PSSC, including one type not previously described. Tailocins were always found to carry only a single tail fiber type, with protein structure predictions of each fiber type suggesting one to be descended from long tail fibers (hereafter type 1a), which in bacteriophage bind reversibly to the host cell, while the other two are descended from short tail fibers (type 2 and 3), which are typically associated with irreversible binding. While we found three distinct tail fiber clades, most strains carried either type 1a or type 2 fibers, which were roughly equally represented in our dataset. Additionally, we found that type 1a tailocin tail fibers are genetically and structurally related to those carried by intact prophage commonly found throughout the species complex. Finally, we found that the distribution of tail fibers throughout PSSC is often incongruent with PSSC phylogeny. Rather, carriage of tail fiber types was highly correlated with the allelic diversity of the LPS gene dTDP-4-dehydrorhamnose reductase (*rfbD*). Our results, when interpreted along with recent findings that tailocin sensitivity also strongly correlates with *rfbD* variation^25^ and the strong similarities in killing spectra exhibited by tail fibers of the same type^21^, suggests the presence of two distinct groups within PSSC, described by their tail fiber and *rfbD* content, that transcend taxonomic classifications, and that are capable of reciprocal killing. These results have implications for the ability of PSSC strains to co-exist with, and suppress the pathogenic potential of, closely related pathogens.

## Methods

### Genomes used in this study

All genomes labeled as ‘*Pseudomonas syringae* species group’ were downloaded on November 17^th^, 2021 from NCBI. A total of 2,468 genomes were checked for completeness and assembly quality with BUSCO, and only genomes with a BUSCO score >= 99 were used in this study (2,161 genomes total).

Phylogroups were assigned to all genomes as outlined in ^27^, with any genomes unassignable to a phylogroup due to sharing less than 95% ANI with any reference strain assumed to be outside PSSC. Although not considered as PSSC members here, these genomes were kept in our dataset, with the logic that they represent the outer boundaries of what might reasonably be considered to be the species group^28,29^.

### Genomic screens for genes of interest

#### Tail fibers

Strains in PSSC are thought to carry and produce a single tailocin, with each tailocin equipped with one of three distinct tail fibers that generally correlate with distinct killing spectra ^21^. A thorough search for all representatives of these three tail fibers within PSSC genomes was conducted using the following method. First, as tailocins in PSSC have been exclusively found encoded in a ca. 12kb locus directly downstream of Anthranilate synthase component 2 (*trpG*) ^11^, we extracted 25 kilobases in this region and annotated tail fiber genes contained within the extraction by first identifying open reading frames with GLIMMER3 ^30^ and then identifying tail fiber genes based on similarity to known tail fiber sequences (Table 1) using the ‘Annotate from Database’ functionality in Geneious 2023.1.1 ^31^. 85% similarity was used a threshold for annotation.

**Table 1.** Tailocin-associated tail fiber reference sequences used to annotate tail fiber genes.

| *PSSC strain* | *Protein accession (NCBI)* |
| --- | --- |
| **Type 1 tail fibers** |  |
| CC1548 | WP_024693886.1 |
| B301D | WP_032656593.1 |
| CC440 | WP_024649699.1 |
| USA007 | WP_024658146.1 |
| **Type 2 tail fibers** |  |
| UB303 | WP_024639976.1 |
| Por 1_6 | WP_005896706.1 |
| CC1583 | WP_024674765.1 |
| CC1630 | WP_005768002.1 |
| **Type 3 tail fibers** |  |
| UB246 | WP_027901668.1, WP_027901669.1 |

Using the tail fiber genes annotated within Geneious, we then screened each genome for tailocin tail fibers that might be located outside of the expected genomic region. We built multiple sequence alignments for each of the three tail fiber types using MAFFT ^32^ under default settings, built HMM profiles with HHMbuild (HMM_1_, HMM_2_, and HMM_3_, Supplementary data 1-3), and screened genomes with HMMscan in HMMER ^33^, with an E-value < 10^−20^ considered to be a significant hit. In instances where a single gene was identified as homologous to multiple tail fiber types, the prediction with the highest E-score was used to assign identity. For tail fiber naming conventions, we strived to stay consistent with both the HMM that best matched the tail fiber and Baltrus et al.’s ^21^ existing numbering scheme for killing classes, such that e.g., tail fibers we designate as type 1 represent those detected by HMM_1_ and also contain fibers in killing class 1.

#### Tailocin- and prophage-associated genes

To help discriminate between tail fibers associated with either tailocins or prophages, we relied on the presence of key genes in the same genomic region as the tail fibers. As mentioned above, tailocins in PSSC have previously only been found directly downstream of *trpG*, with *trpE* and *trpD* downstream of the tailocin region ^11^. Therefore, we interpreted the presence of these genes near the tail fibers to suggest that the tail fibers are associated with a tailocin. Additionally, while prophages require terminase and capsid genes for packaging and storing DNA ^34^, tailocins lack these genes^35^. Thus, the presence of capsid and terminase genes suggests that any tail fibers nearby are associated with a prophage.

Genomic CoDing Sequences (CDS) for each genome were downloaded from NCBI. Using an in-house script, 25 CDS upstream and downstream of each tail fiber identified above were searched for presence of the terms ‘*trpG*’, ‘*trpE*’, ‘Anthranilate synthase component II’, ‘capsid’, and ‘terminase’ in their gene names and descriptions. 25 CDS was found to be a conservative threshold for capturing the entire tailocin locus considering a typical PSSC tailocin has 27 CDS^11^. As only a small proportion of genomes in our dataset were fully circularized, it was expected that some of our indicator genes would not be on the same contig as the focal tail fiber, leading to false negatives. Therefore, presence of any of the above genes were considered evidence of a tail fiber belonging to the appropriate particle type (tailocin or prophage). In ambiguous cases, e.g. when both prophage and tailocin-associated genes were present near a tail fiber, or no diagnostic genes were found, manual inspection of the loci, the phage search annotation web tool PHASTER^36^, and the phylogenetic relationship to other fibers were considered when assigning an identity.

#### rfbD genes

Using the method described above, all CDS annotated as ‘*rfbD*’ were extracted from the PSCC genome set.

### Phylogenetic trees

Gene trees for tail fibers detected by each HMM used in this study were generated by first reducing each set of genes to non-redundant sequences (i.e. sequences with unique accession numbers). Amino acid sequences were then aligned using MAFFT (ref for MAFFT) under default settings, an alignment mask removing columns containing greater than 20% gaps was applied, and a phylogenetic tree was built using a Jukes-Cantor distance model and the neighbor-joining method. Both the alignment mask and phylogenetic tree were implemented with Geneious Prime 2023.1.1 ^31^.

As the *rfbD* analysis presented was concerned with PSSC genomes only, any genomes for which we were unable to assign phylogroups were removed from the analysis. A gene tree for *rfbD* was built using the same method used to build the tail fiber gene trees above. *rfbD* genes found in all three genomes within phylogroup 13, which act as outgroup to all other phylogroups within PSSC, were used as an outgroup and subsequently removed from the tree for clarity in the final figure.

### AlphaFold protein structure prediction

Tail fibers equipped by both bacteriophage and tailocins are frequently homotrimeric ^12,13,37^. In accordance with this, we present predicted structures of all tail fibers as homotrimers.

Multimers were predicted with AlphaFold 2.3.0 ^38^, and the top ranked relaxed models were chosen for inclusion in the final manuscript. All predicted structures and amino acid sequences used for prediction are available in Supplementary data 3-13.

## Results and Discussion

### Diversity of tailocin-associated tail fibers within PSSC

Members of PSSC are commonly categorized into phylogroups, with a distinction between primary phylogroups (phylogroups1,2,3,4,5,6 and 10) ^39^ that are closely related to each other, and the more distant secondary phylogroups (7, 9, 11, and 13). While the secondary phylogroups do contain pathogens, most notably *P. viridiflava* from phylogroup 7 ^40^, the vast majority of highly virulent pathogens infecting common crops are found in the primary phylogroups ^41^. Among all genomes we surveyed, 62% were found to be carrying a tailocin (differentiated from prophages by a lack of terminase and capsid genes) directly downstream of the *trpG* gene, with no evidence of tailocins elsewhere in any PSSC strains. Tailocins were most common within the primary phylogroups (1,2,3,4,5,6 and 10), in which the proportion of genomes carrying tailocins increased to 84%. As the primary phylogroups are most highly represented in our dataset, contain the most agriculturally significant pathogens ^39^, and also carry tailocins most frequently, we decided to focus primarily on this group.

We screened genomes for three previously described tail fiber types that are differentiated both by large recombination events in the coding sequences, and a marked difference in their killing spectra ^21^. HMMs discussed here and subscripted with 1, 2, and 3 therefore correspond to three distinct clades of tail fibers described by Baltrus et al. ^21^. HMM_1_ and HMM_2_ returned 1,619 and 306 significant hits representing 290 and 63 nonredundant amino acid sequences, respectively.

Tailocin tail fibers found by HMM_1_ form two monophyletic clades (Figure 1a), one of which contains fibers found exclusively in Phylogroup 7, and the other which contains those found throughout the primary phylogroups 1,2,3,5, and 6. Although these two clades of fibers are distinct evolutionarily, they appear to share a common ancestor, as they share strong homology throughout the length of the tail fibers (see pairwise alignment of type 1a and 1b, Figure 1b). Therefore, HMM_1_ fibers associated with tailocins and found in the primary phylogroups and Phylogroup 7 are hereafter referred to as Type 1a and 1b, respectively.

**Figure 1.**
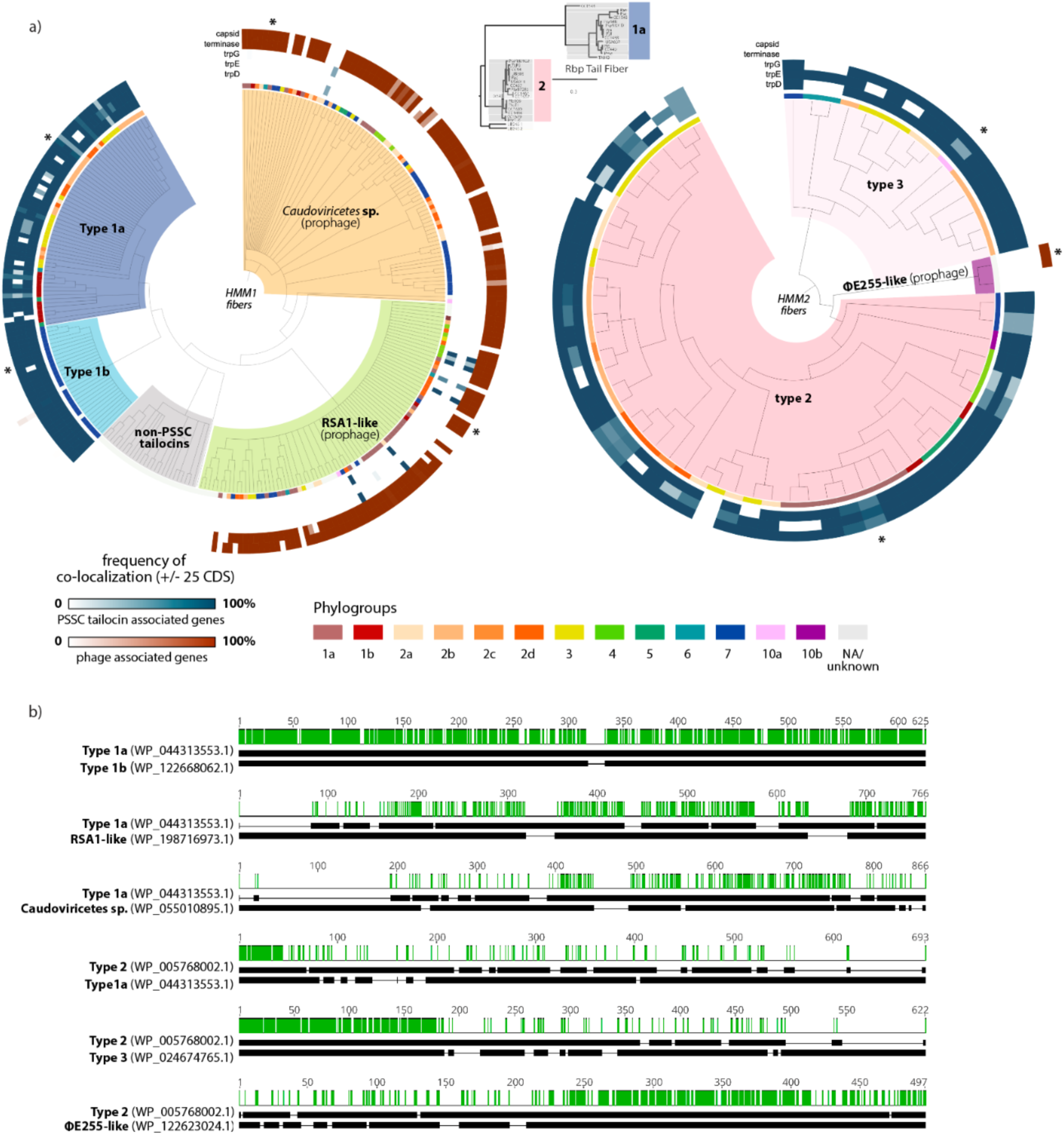
Tail fibers associated with PSSC tailocins exhibit more diversity than previously reported and are closely related to prophage fibers found in PSSC genomes. **a)** gene trees for non-redundant tail fiber sequences detected by HMM_1_ (left) and HMM_2_ (right). Dendrograms are colored and labeled according to tail fiber types. Annotation rings from the inside out: 1) Phylogroup of the genomes containing each tail fiber; 2-4) blue heatmaps show the frequency of *trpD*, *trpE*, and *trpG* being detected within 25 CDS of the tail fiber; 5,6) red heatmaps show the frequency of terminase and capsid genes being detected within 25 CDS of the tail fiber. Asterisks denote tail fibers used for pairwise alignments in panel b. The central rectangular tree is a modified tail fiber gene tree from ^21^. Color and number annotations to the right of the tree correspond with the naming conventions used for fiber types in this study. **b)** pairwise amino acid sequence alignments for representative tail fibers, with green indicating conserved residues. Tail fiber group that the sequences are representing, along with the non-redundant protein accession numbers are provided to the left of each alignment.

Tail fibers identified with HMM_2_ were comprised of fibers closely related to those associated with killing class 2, with a small subclade showing evidence of a large recombination event replacing the latter half of the gene (see pairwise alignment between type 2 and type 3, Figure 1b). A representative fiber from this clade (found in strain CC1583) was among those investigated for killing activity by Baltrus et al. ^21^, but was dismissed as a truncated Type 2 fiber in the phylogenetic analysis. Nonetheless, CC1583’s tailocin appeared to be functional and was described as belonging to killing class 2. Despite the similarity in the killing spectra that has been reported, due to the marked genetic and structural differences between this clade and the rest of the type 2 fibers, we hereafter refer to this clade of tail fibers as Type 3 (Figure 1a).

Tail fibers identified with HMM_3_ were almost exclusively found in genomes outside of PSSC and thus further analysis of the tail fibers was considered beyond the scope of this study.

Supplementary data 14 describes the full tail fiber contents of genomes screened in this study.

Fibers identified by HMM_3_ were found predominately in *Pseudomonas* strains outside any currently described phylogroup (Supplementary data 15), with thirteen instances in secondary phylogroups, and only a single instance within the primary phylogroups (ICMP-19589, phylogroup 3). It was determined that a proper accounting of the fibers detected by HMM_3_ would require a broader screening of tailocins throughout the *Pseudomonas* genus, and further investigation of these fibers was deemed to be beyond the scope of this paper.

Tailocin fibers detected in PSSC (Type 1a, 1b, 2, and 3) all share strong homology of at least 60 n-terminal amino acid residues (Figure 1b), with total protein lengths ranging from ca. 400-800 residues. The conserved n-terminal region is thought to be important for attachment of phage tail fibers to the baseplate ^21,37,42^, and it is possible that 60 amino acids represents a minimally required domain for proper assembly and/or functioning of tail fibers with the PSSC tailocin. This is potentially significant for any future attempts to engineer synthetic versions of the PSSC tailocins, as it suggests the ability to fuse novel binding domains to the conserved n-terminal region to retarget killing spectra beyond any wild-type capabilities of the tailocin, as has been done with tailocins produced by *P. aeruginosa* ^14,26^.

### Extant prophages within PSSC genomes carry tail fibers closely related to Type 1 tailocin fibers

Our genomic screen revealed more diversity in tail fibers than previously described; several genomes carried more than one tail fiber, despite inspection of tailocin regions in PSSC consistently revealing them to only carry a single fiber. We considered two possible explanations for this observation: 1) Additional tailocins were present in PSSC, or 2) there are prophages within PSSC that are equipped with tail fibers closely related to known tailocin tail fibers. In an attempt to tease apart these two hypotheses, we screened the genomic region flanking tail fiber genes looking for diagnostic genes that would indicate that the tail fiber belongs to a tailocin or prophage. Specifically, as the PSSC tailocin of interest is thought to always be located immediately adjacent to the trp operon, we looked for the presence of *trpG*, *trpE*, and *trpD* within +/-25 ORFs of all tail fibers to indicate the fibers as being associated with the tailocin. Additionally, to assess whether any fibers might belong to prophages, we also searched this region for capsid and terminase genes – which are essential for tailed bacteriophages, but not for tailocins ^43^. We found that many tail fibers closely related to type 1 tailocins were adjacent to capsid and terminase genes, but none of the trp genes (red and blue annotation rings, Figure 1a).

Further analysis of the putative prophages with PHASTER suggested these are indeed intact prophages. As an illustrative example, the analysis of such a region found in *P. syringae* pv. tomato strain Pst-DC-98-1 being identified as an intact phage with a score of 91 (scores >90 are considered to be indicative of an intact phage by PHASTER), with 22/38 identified phage genes being most homologous to *Pseudomonas* phage <13. Specific identity varies considerably between phages characterized with PHASTER, particularly when focusing on the tail fiber gene. Therefore, for practical purposes, we describe the origin of most of these fibers as simply belonging to the tailed bacteriophages, or *Caudoviricetes* phages (Figure 1a). An exception to this is a monophyletic clade of fibers that consistently BLAST as being closely related to RSA1 tail fibers, and thus this clade is described as RSA1-like (Figure 1a). RSA1-like fibers exhibit greater homology to both type 1a and 1b tailocin fibers than the aforementioned *Caudoviricetes* sp. fibers in all but the first 100 residues, although large deletions between RSA-1 like fibers and type 1 fibers are common (Figure 1b).

Nearly all fibers identified by HMM_2_ were found to be associated with tailocins. Two putative prophages carrying partially homologous fibers were, however, found in two genomes – *Pseudomonas petrae* (GCF_900585705.1) and an additional strain (GCF_001698815.1) whose identity is unclear, but shares 84% average nucleotide identity with *Pseudomonas foliumensis,* according to NCBI’s taxonomy check. The regions surrounding these putative phage fibers were also determined by PHASTER to be intact prophages (score 140), with the fibers most closely related to phage <1E255 (<1E255-like, Figure 1a). Homology between <1E255-like fibers and Type 2 fibers was restricted to the c-terminal region (Figure 1b). Given the consistent pattern of highly similar c-terminal regions being found in circulating *Pseudomonas* phages, it is tempting to speculate that these phages were the tail fiber donors for type 1 and 2 c-terminal regions seen in PSSC tailocin fibers, although it is difficult if not impossible to confirm this.

### Structural comparison of tail fibers and their functional implications

Given the extensive recombination evident among the tail fibers described above, we sought to better understand the impact that such genetic diversity has on protein structure. We used AlphaFold to predict structures for representatives from each clade of fibers and found that overall, type 2 and 3 tail fibers likely derive from short tail fibers while type 1a and 1b are likely long tail fibers with structural similarities to the *P. aeruginosa* R2 tailocin tail fiber ^42^. All fibers were modeled as homotrimers, in line with observations from various phage and tailocin tail fibers ^12,44^.

#### Type 1: the long tail fibers

The prototypical type 1 tail fiber is approximately 450 AA long, with an n-terminal baseplate attachment point, followed immediately by a large knob domain of unknown function. The majority of the tail fiber is comprised of three repeated units (Figure 2a), with each repeat consisting of a shaft, an n-proximal knob (knob a) and a c-proximal knob (knob b) (Figure 2a). These repeats are not just structurally similar but share strong homology at the amino acid level across the length of the repeat (Figure 2b).

**Figure 2.**
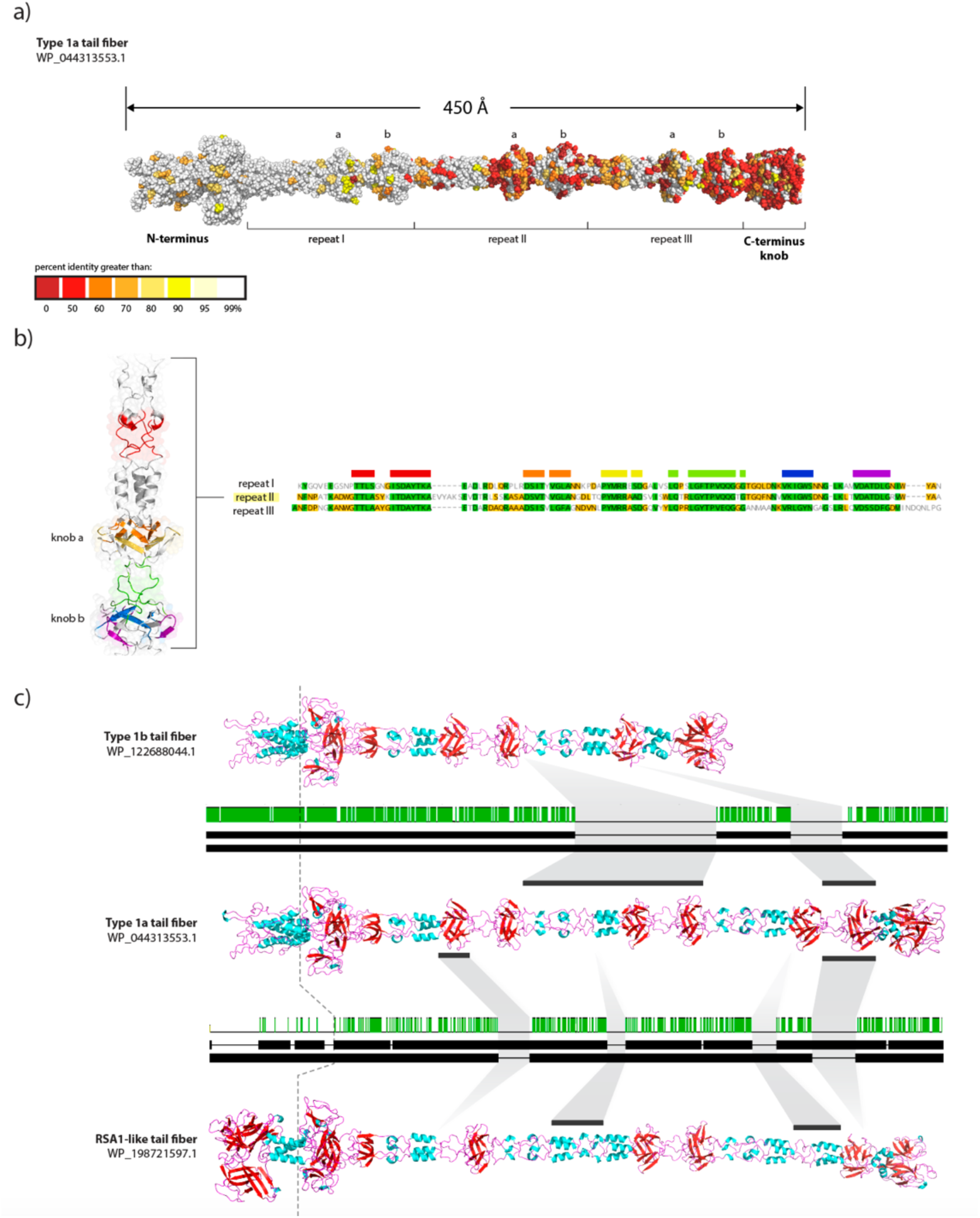
Type 1 tail fibers and their phage relative display modular structure. **a)** AlphaFold-predicted structure of a typical type 1a tail fiber, colored by conservation of residues. The n-terminus baseplate attachment point is far left, with Repeats 1, 2, and 3 labeled below the tail fiber. Knob domains a and b within each repeat are labeled above tail fiber. **b)** Amino acid multiple sequence alignment for repeats 1, 2, and 3 from the tail fiber in panel a. Cartoon ribbon structure for repeat 2 is to the left, colored in accordance with the highly conserved regions seen in the MSA. **c**) AlphaFold-predicted structures for tail fibers belonging to type 1b, 1a, and the RSA1-like phage. Between each pair of fibers is shown the accompanying amino acid pairwise alignment. Dotted line running through all fibers to the left denotes the large structural difference in the baseplate-attachment domain between tailocin-associated fibers (type 1a and 1b) and the RSA1-like fibers. The light grey annotations between fibers highlight large deletions present in some fibers tend to center around whole repeats or single knob domains.

The general architecture of type 1 tailocin fibers bears resembles the crystal structures for the c-terminal regions of R1 and R2 pyocin tail fibers, which contain a single pair of knobs followed by a c-terminus knob ^37^. Each of these knobs have been suggested to function as receptor binding domains, providing some evidence for the role of these structures in binding for type 1 fibers – although type 1 tail fibers and the R2 pyocin tail fiber from *P. aeruginosa* PA01 (PA0620) bare little similarity to each other at the amino acid level (Supplementary Figure 1).

Among type 1a fibers, amino acid conservation was the highest in the n-terminal attachment domain, and decreased steadily over the length of the fiber, with areas of low percent identity concentrated primarily in knobs found in repeats II and III, as well as the c-terminal knob (fig. 2a). These results suggest that the distal binding domains might play a greater role in attachment to target cells than proximal knobs, and thus are under greater selection.

The general architecture between type 1a, 1b, and RSA1-like fibers is strikingly similar, save for large deletions that commonly encompass entire knob domains. In an example of typical deletions found among tail fibers is highlighted in figure 2c, in which repeat II was deleted entirely in a small group of type 1b fibers, along with knob b in repeat III. The result is a significantly shorter tail fiber with three internal knobs instead of the typical six. Also shown in figure 2c is an instance where knob a in repeat I as well as knob b in repeat III of an RSA1-like fiber are missing, when compared to the prototypical type 1a fiber. The frequency of such large deletions (multiple sequence alignment of all type 1 tail fibers, supplementary Figures 2 and 3) strongly suggests there is significant functional importance for the predicted knob domains, and deletions of individual knobs might play a role in shaping killing spectra associated with a given tail fiber, although experimental work would be needed to determine what if any role these deletions play.

**Figure 3.**
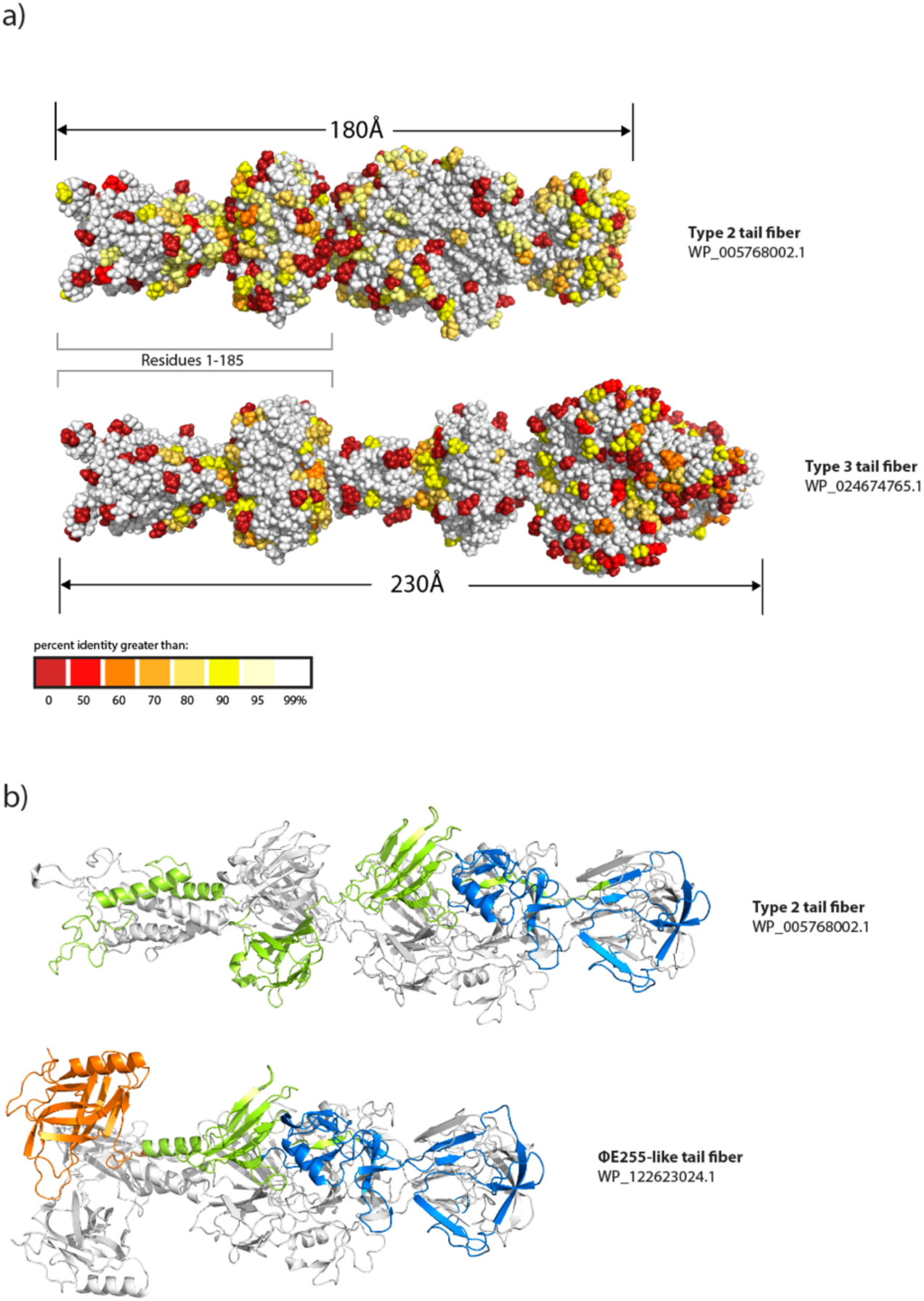
Structural similarities between type 2, 3, and <ΔE255-like fibers, as predicted by AlphaFold. **a)** Structures of representative fibers belonging to type 2 and type 3, colored by residue conservation. The highly conserved 185 residue region seen in fig. 1b is highlighted with the open rectangles. **b)** Structures of a type 2 and <ΔE255-like fiber are shown in ribbon form, with chain b and chain c colored white, and chain a colored to highlight the DUF3751 domain (orange), T4 short tail fiber binding domain (blue), and regions with no known function (green) according to InterProScan.

As noted above, the n-terminal region of RSA1-like fibers share no homology with tailocin-associated fibers (Figure 1b), and the predicted protein structures recapitulate these differences by exhibiting significant structural differences in this region (residues 1-156) compared to the otherwise similar type 1a fibers (Figure 2c). Functionally, this is a potentially significant difference, as it suggests that despite otherwise strong genetic and structural similarity, it is unlikely that RSA1-like prophage fibers are compatible with PSSC tailocin particles (i.e., a PSSC strain carrying both a tailocin and such a prophage is unlikely to be able to incorporate the prophage fibers into the structure of the tailocin).

#### Types 2 and 3: the short tail fibers

Structures of type 2 and 3 tail fibers are shorter and more globular than type 1 fibers (Figure 3a), suggesting both derive from short tail fibers. Reflecting the pattern of conservation seen in their amino acid sequences, a large segment (185AA) comprising the conserved baseplate attachment domain and a large knob of unknown function is shared in both fibers. The distal half of the fibers share no discernable similarities, and neither fiber appears to have any regions with particularly high sequence diversity, as opposed to type 1 fibers.

Sequences for both tail fiber types were scanned for known protein domains with InterProScan, and while no conserved domains were detected in the type 3 fibers, a large region in the distal half of the type 2 fibers was found to be homologous to the T4 short tail fiber binding domain (Figure 3b). This domain was also found in the <1E255-like fibers, as was a DUF3751 domain at the baseplate attachment point of the fiber, which have been associated with both prophage and tailocin fibers in *P*. *putida* ^10^.

The significance of PSSC tailocins carrying either short or long tail fibers is unclear. In the infection cycle of bacteriophage, both of these fibers play distinct roles, with long tail fibers binding reversibly to the cell surface, allowing the phage to ‘walk’ around until a sufficient number of fibers have bound to trigger a conformational change in the baseplate ^12^. Following the conformational change, the phage lowers onto the cell surface and short tail fibers bind irreversibly to the target cell, allowing more efficient infection ^12^. Whether the nature of reversible vs. irreversible binding remains in tailocin fibers, or if these differences alter the killing efficiency of the particle remains untested.

### Distribution of Type 1 and 2 fibers is highly correlated with LPS gene *rfbD*

Having described the richness of tail fiber types carried by PSSC tailocins, we sought to better understand their distribution and relative abundance among the genomes in our dataset. Among the primary phylogroups, type 1a and type 2 tail fibers were fairly evenly represented, at 38% and 42%, respectively, with only 4% of genomes carrying type 3 fibers. Surprisingly, however, the proportion of each fiber type varied greatly by phylogroup. For phylogroups 2, 3, and 6, which formed a monophyletic group in our core genome tree, both type 1a and 2 fibers were again evenly represented (table 2). However, phylogroups 1a, 1b, 4, and 5, which also formed a monophyletic group, tended to be dominated by one fiber type or the other (table 2). If these distributions represent genuine differences in tail fiber abundances between clades in PSSC, as opposed to being the result of sampling bias in some way, it suggests that perhaps tail fibers are under more heightened selection and undergo more frequent recombination in some phylogroups than others. For instance, the most virulent pathogens tend to reside in phylogroup 1, whereas phylogroup 2 is considered to contain strains that are more widespread, being better epiphytes and generally less virulent ^41^. In this instance, tailocins might provide a greater fitness advantage to strains that spend more time on the leaf surface than those that spend more time isolated from other microbial competitors inside the plant apoplast. However, it is also possible that sampling biases of highly clonal aggressive pathogens skew this dataset, leading to an underestimation of the true diversity of tail fibers among phylogroups 1a, 1b, 4, and 5.

**Table 2.** Proportion of genomes carrying each tailocin-associated tail fiber. PG = phylogroup, n = # of genomes.

| PG (n) | Type 1a | Type 1b | Type 2 | Type 3 |
| --- | --- | --- | --- | --- |
| 1a (108) | 0 | 0 | 0.92 | 0 |
| 1b (137) | 0.93 | 0 | 0.04 | 0 |
| 2a (36) | 0.11 | 0 | 0.41 | 0.08 |
| 2b (118) | 0.40 | 0 | 0.32 | 0.11 |
| 2c (8) | 0.5 | 0 | 0.5 | 0 |
| 2d (40) | 0.48 | 0 | 0.52 | 0 |
| 3 (125) | 0.352 | 0 | 0.296 | 0.04 |
| 4 (44) | 0 | 0 | 1 | 0 |
| 5 (9) | 0.33 | 0 | 0.66 | 0 |
| 6 (9) | 0.44 | 0 | 0 | 0.22 |
| 7 (1332) | 0 | 0.589 | 0.002 | 0.003 |
| 9 (2) | 0 | 0 | 0 | 0 |
| 10a (2) | 0 | 0 | 0 | 1 |
| 10b (2) | 0 | 0 | 1 | 0 |
| 11 (12) | 0 | 0 | 0 | 0 |
| 13 (3) | 0 | 0 | 0 | 0 |

We also looked at the co-distribution of tail fibers associated with tailocins and prophages (Figure 4a). We found a very strong correlation between genomes carrying tailocins equipped with type 2 fibers and those harboring prophages. Among genomes carrying type 2 fibers, 54% also carried at least one prophage fiber. In contrast, among genomes carrying type 1a fibers, only 4% of genomes carried any prophage fiber. Considering the structural similarities between the prophage and type 1a fibers established above, and assuming structural similarities correspond with similar binding affinities to LPS motifs, this distribution pattern suggests that PSSC strains that are able to be infected by phages equipped with the RSA-1 like tail fibers might also be less likely to carry a type 1a tail fiber with their tailocin due to an increased chance of self-killing.

**Figure 4.**
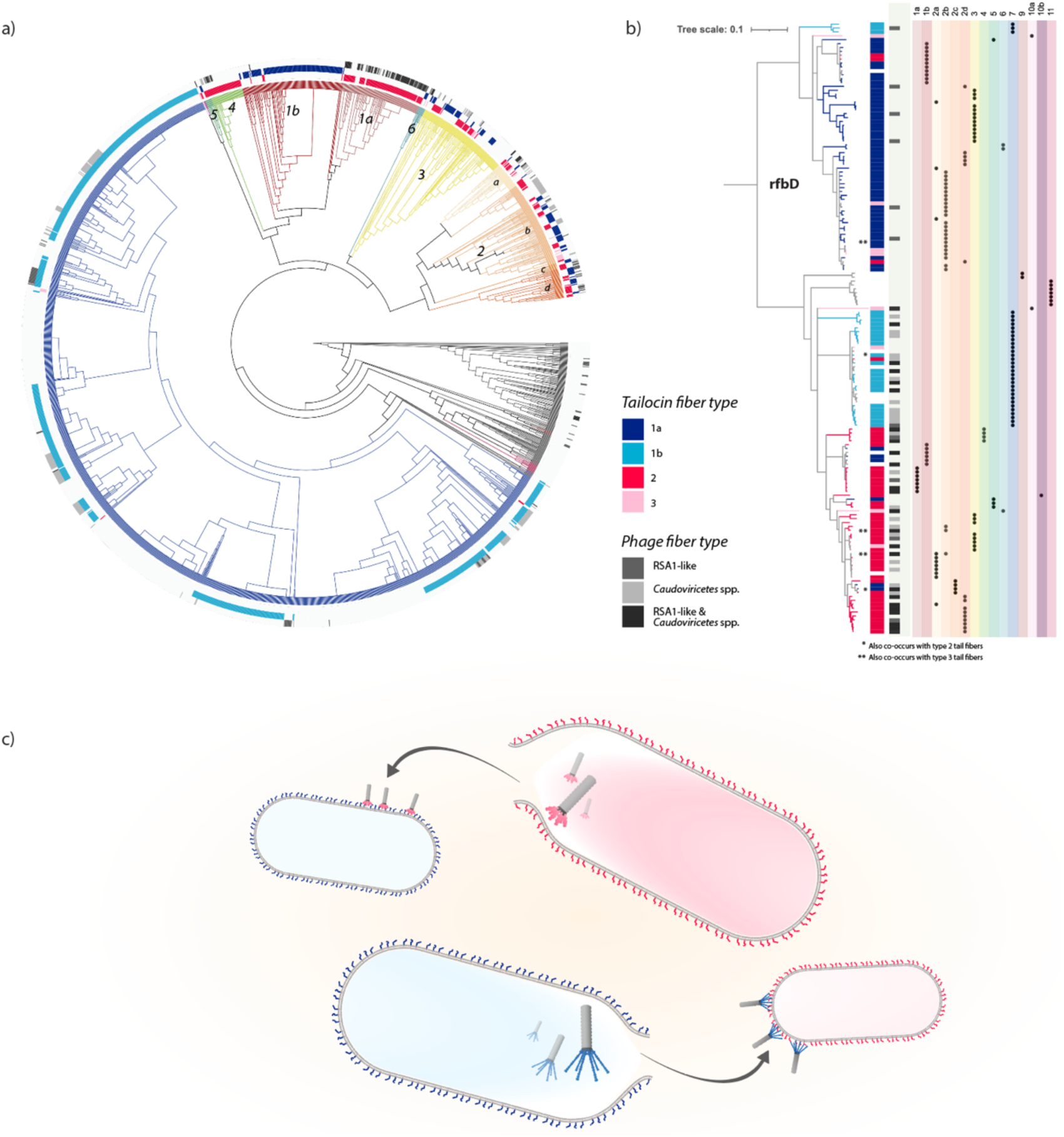
Distribution of tail fiber types correspond better to presence of *rfbD* alleles than to phylogroup. **a)** core genome phylogeny of PSSC, with branches colored according to phylogroup. Primary phylogroups are numbered. Grey branches are genomes that were too distant from reference strains to confidently assign to any known phylogroup. Three annotations rings represent, from inside to out, 1) presence of type 2 (dark pink) and 3 (light pink) tail fibers, 2) presence of type 1a (dark blue) and 1b (light blue) fibers, and 3) presence of phage fibers belonging to a broad group of *Caudoviricetes* phage (light grey), RSA1-like phages (mid-grey), or both (dark grey). **b)** a gene tree for *rfbD*, rooted at alleles found exclusively in phylogroup 13. Tree branches and the inner annotation strip are colored according to the tailocin-associated tail fiber types that were observed to be co-occurring in the same genome with each *rfbD* gene sequence. Instances where an *rfbD* sequence is seen co-occurring with multiple tail fiber types are marked with asterisks. The greyscale strip represents *rfbD* co-occurrences with phage-associated tail fibers. To the right, the phylogroup of origin for each *rfbD* sequence is shown. **C)** Conceptual model for the hypothesis that LPS structure, as differentially conferred by *rfbD* allele, resulting in the reciprocal killing of PSSC strains in the species complex.

It has recently been reported that sensitivity to tailocins in PSSC is strongly linked to the carriage of LPS-related genes, and that allelic variation in *rfbD* is a particularly robust predictor ^25^. Given that *rfbD* is a strong predicter of tailocin sensitivity, we hypothesized that it might also reflect a barrier to carriage of certain tail fibers, as to avoid self-killing. Additionally, because we observed two dominant tail fiber types (type 1a and 2) and *rfbD* alleles within PSSC form two distinct clades associated with tailocin sensitivities, we hypothesized that the allele of *rfbD* carried by a genome would be strongly correlated with the type of tail fiber carried. Finally, we hypothesized that due to the significant structural similarities between type 1a and 1b fibers, the co-distribution pattern between type 1a fibers and *rfbD* alleles would resemble the pattern between type 1b fibers and *rfbD* alleles. To test these hypotheses, we built a gene tree of *rfbD*, and analyzed co-occurrence with each fiber type (1a, 1b, 2, and 3). For the sake of completeness, we also investigated the co-occurrence of *rfbD* alleles with RSA1-like and *Caudoviricetes* fibers.

Consistent with the results by Baltrus et al. ^25^, two major clades of *rfbD* were observed. Each clade exhibited distinct correlations with both tailocin-associated and prophage-associated fibers (Figure 4b). Within the first clade of *rfbD* genes, 84% of unique alleles were associated with type 1a fibers, only 9% were associated with any prophage fiber, and 5% were associated with type 2 fibers. In contrast, in the second clade of *rfbD* genes, 7% of unique alleles were associated with type 1a fibers, 58% were associated with at least one prophage-associated fiber, and 49% were associated with type 2 fibers. These results are consistent with the hypothesis that the distribution of tail fibers observed in PSSC is determined at least in part by aspects of LPS structure.

A surprising result from the analysis of tail fiber co-occurrence with *rfbD* alleles is that despite being both genetically and structurally very similar to type 1a tail fibers, type 1b fibers are associated with *rfbD* alleles that are much more like those commonly found in strains carrying type 2 tail fibers than with type 1a fibers. Assuming that it is necessary for a tailocin-producing strain to carry a tail fiber that can co-exist with its rhamnose-synthesizing gene *rfbD*, this result implies that despite the similarity between type 1a and 1b fibers, either their binding activity and therefore killing spectra are distinct, or that despite phylogenetic similarity between *rfbD* alleles found in genomes with type 2 and type 1b fibers, the LPS they ultimately produce are structurally distinct. Whichever is the case, the interplay between *rfbD* and tailocin activity warrants further investigation.

Finally, it is worth commenting on the significance of the observed correlation between extant prophages in PSSC and alleles of *rfbD*. While it has been well documented that many *Pseudomonas* phages target LPS ^22,45^, and that these phages exhibit differential killing spectra ^46,47^, to our knowledge we are presenting here the first evidence for a simple mechanism that might be driving phage-bacteria dynamics within PSSC: compatibility with one of two *rfbD*-dependent LPS structures that decorate PSSC outer membranes. If, like tailocins, PSSC phage populations turn out to be heavily biased toward certain strains based on the activity of *rfbD* within host strains, the implications for the evolution of PSSC could be significant, as phages have been implicated in the horizontal gene transfer of effector proteins between PSSC strains ^48^. From the perspective of management, the results here might also provide insight for more thoughtful production of phage therapy cocktails ^49^

## Supplementary Figures

**Supplemental Figure 1.**
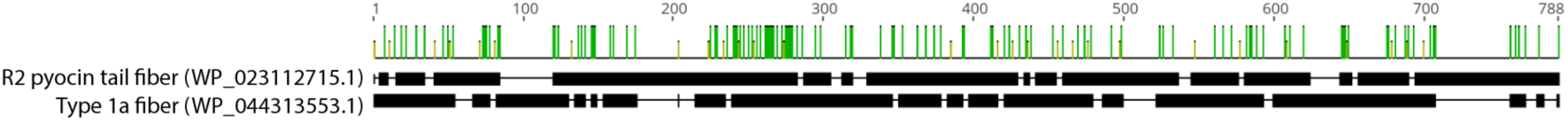
Pairwise alignment of type 1a tailocin-associated tail fiber found in PSSC and an R2 pyocin fiber from *P. aeruginosa*

**Supplemental Figure 2.**
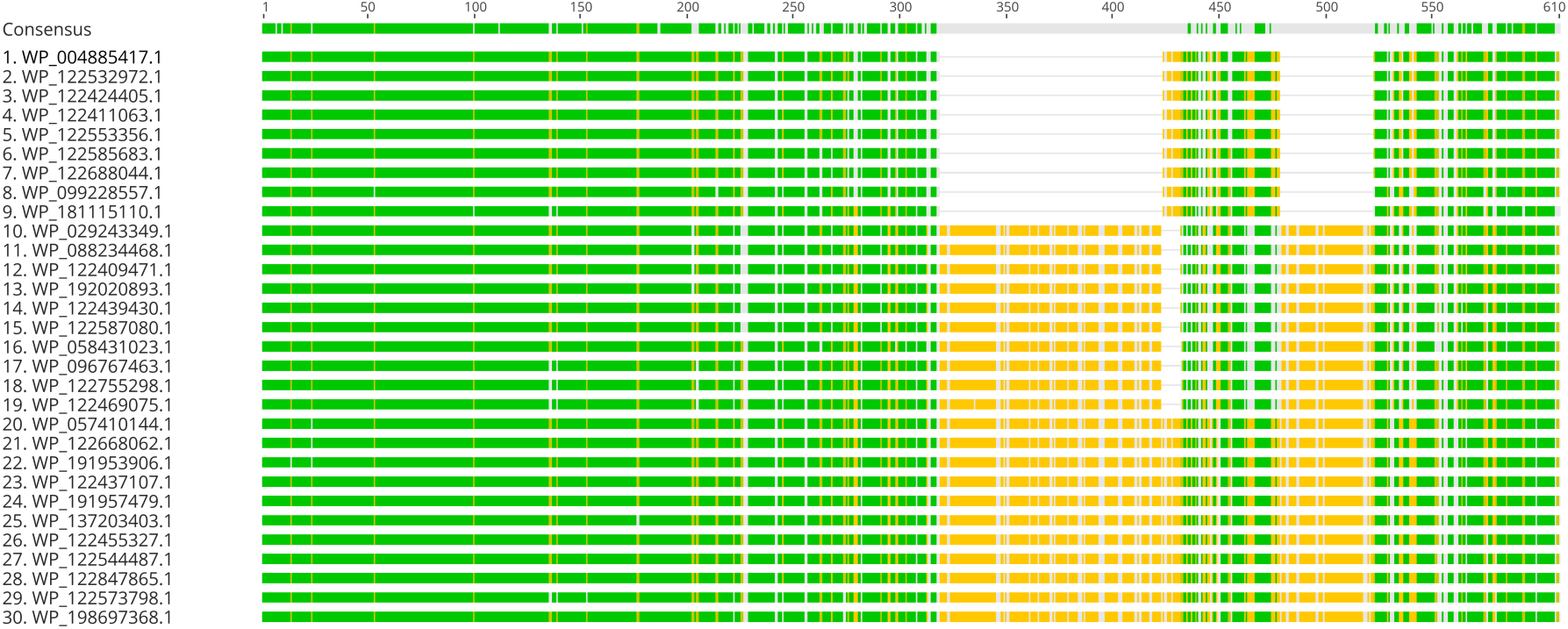
Multiple amino acid sequence alignment for all unique type 1b fibers found in this study

**Supplemental Figure 3.**
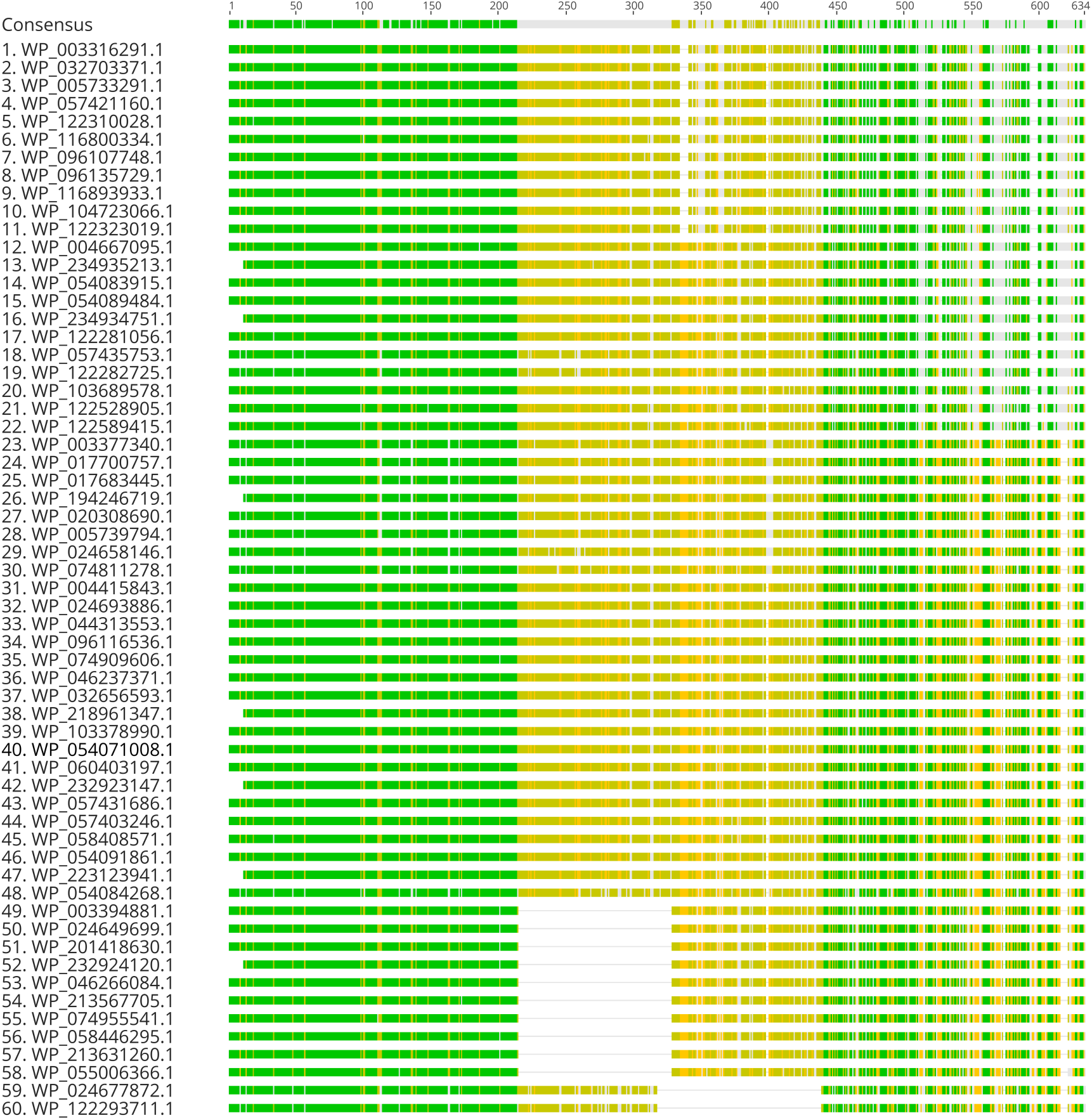
Multiple amino acid sequence alignment for all unique type 1a fibers found in this study

## Supplementary data

All supplementary data are deposited at Zenodo ^50^. Data include:

### Supplementary data 1

HMM_1_, representing tailocin tail fibers associated with killing class 1

### Supplementary data 2

HMM_2_, representing tailocin tail fibers associated with killing class 2

### Supplementary data 3

HMM_3_, representing tailocin tail fibers from PSSC strain UB246

### Supplementary data 4

Amino acid sequence for WP_044313553.1, representative of type 1a tailocin-associated tail fiber used for protein structure prediction in Supplementary data 3.5

### Supplementary data 5

PDB file containing predicted structure of WP_044313553.1, representative of type 1a tailocin-associated tail fiber

### Supplementary data 6

Amino acid sequence for WP_122688044.1, representative of type 1b tailocin-associated tail fiber used for protein structure prediction in Supplementary data 3.7

### Supplementary data 7

PDB file containing predicted structure of WP_122688044.1, representative of type 1b tailocin-associated tail fiber

### Supplementary data 8

Amino acid sequence for WP_005768002.1, representative of type 2 tailocin-associated tail fiber used for protein structure prediction in Supplementary data 3.9

### Supplementary data 9

PDB file containing predicted structure of WP_005768002.1, representative of type 2 tailocin-associated tail fiber

### Supplementary data 10

Amino acid sequence for WP_024674765.1, representative of type 3 tailocin-associated tail fiber used for protein structure prediction in Supplementary data 311

### Supplementary data 11

PDB file containing predicted structure of WP_024674765.1, representative of type 3 tailocin-associated tail fiber

### Supplementary data 12

Amino acid sequence for WP_198721597.1, representative of RSA1-like prophage-associated tail fiber used for protein structure prediction in Supplementary data 3.13

### Supplementary data 13

PDB file containing predicted structure of WP_198721597.1, representative of RSA1-like prophage-associated tail fiber

### Supplementary data 14

CSV file containing HMM_1_ and HMM_2_ genomic screen results, with accession numbers and identities of tail fibers detected in each genome.

### Supplementary data 15

CSV file containing HMM_3_ genomic screen results, with copy number of tail fibers detected in each genome and the phylogroup the genome belongs to

